# Deleterious functional consequences of perfluoroalkyl substances accumulation into the myelin sheath

**DOI:** 10.1101/2023.05.30.542807

**Authors:** L. Butruille, P. Jubin, E. Martin, MS. Aigrot, M. Lhomme, JB. Fini, B. Demeneix, B. Stankoff, C. Lubetzki, B. Zalc, S. Remaud

## Abstract

Exposure to persistent organic pollutants during the perinatal period is of particular concern because of the potential increased risk of neurological disorders in adulthood. Here we questioned whether exposure to perfluorooctanoic acid (PFOA) and perfluorooctane sulfonate (PFOS) could alter myelin formation and regeneration. First, we show that PFOS, and to a lesser extent PFOA, accumulated into the myelin sheath of postnatal day 21 (p21) mice, whose mothers were exposed to either PFOA or PFOS (20mg/L) *via* drinking water during late gestation and lactation, suggesting that accumulation of PFOS into the myelin could interfere with myelin formation and function. In fact, PFOS, but not PFOA, disrupted the generation of oligodendrocytes, the myelin-forming cells of the central nervous system, derived from neural stem cells localised in the subventricular zone of p21 exposed animals. Then, cerebellar slices were transiently demyelinated using lysophosphatidylcholine and remyelination was quantified in the presence of either PFOA or PFOS. Only PFOS impaired remyelination, a deleterious effect rescued by adding thyroid hormone (TH). Similarly to our observation in the mouse, we also showed that PFOS altered remyelination in *Xenopus laevis* using the Tg(*Mbp:GFP-ntr*) model of conditional demyelination and measuring, then, the number of oligodendrocytes. The functional consequences of PFOS-impaired remyelination were shown by its effects using a battery of behavioural tests. In sum, our data demonstrate that perinatal PFOS exposure disrupts oligodendrogenesis and myelin function through modulation of TH action. PFOS exposure may exacerbate genetic and environmental susceptibilities underlying myelin disorders, the most frequent being multiple sclerosis.

**Highlights:**

- Our investigation points the deleterious effects of PFOS incorporation into the myelin sheath
- PFOS interfere dramatically with the generation of remyelinating and functional repair of demyelinating lesions
- Our study points to a potential link between these persistent pollutants and the recent increase in prevalence of multiple sclerosis

## 1. Introduction

Perfluoroalkyl substances (PFAS) are a set of synthetic man-made chemicals produced for over 60 years and widely found in biota, including humans (Calafat et al., 2019; Houde et al., 2016; Stubleski et al., 2017).The most common of these commercial products - largely detected in cosmetics, textiles, food packaging - are perfluorooctanesulfonic acid (PFOS) and perfluorooctanoic acid (PFOA). Due to their long-fluorinated alkyl chain, the half-life of PFAS is exceptionally long (Li et al., 2018). Thus, they are particularly persistent in the environment, longer than any other environmental substance, especially in the human body for which the half-life is estimated about 4 years for PFOA and PFOS (Olsen et al., 2017; Sunderland et al., 2019; Zhang et al., 2013). Several epidemiological studies and experimental analysis using animal models have demonstrated health consequences of PFOA and PFOS including alterations of lipid metabolism, hepatotoxicity, reproduction function as well as impacts on development since PFAS can cross placenta and accumulate in breast milk, thus reaching the offspring (Caporale et al., 2022; Demeneix, 2014). However, there is a paucity of experimental studies evaluating neurobehavioral and molecular mechanisms of neurotoxicity for PFAS (Starnes et al., 2022). PFOA and PFOS have been regulated to limit their use under the Stockholm Convention. However, the current bioaccumulation of these PFAS is so high, that a better understanding of the mechanisms by which PFAS could alter homeostasis is still crucial. Moreover, PFAS are amphiphile compounds that could stick to surfaces and accumulate in adipose tissue (Lee et al., 2017) and potentially in other lipid-rich structures, such as myelin sheath, the oligodendroglial membrane wrapped around long projecting axons. Neurodevelopment and especially the generation of myelin-generating cells are controlled by thyroid hormones (THs). Many chemicals, including PFOS and PFOA, through their TH disrupting effects could induce adverse effects on brain development, thus leading to neurodevelopmental disorders. In particular, the generation of mature myelinating oligodendrocytes depends on TH throughout life, from early development to adulthood. Previous work in the adult mouse demonstrated that a transient lack of TH enhanced the generation of oligodendrocyte precursors (OPCs), derived from neural stem cells (NSCs) within the murine subventricular zone (SVZ). Moreover, these newly-generated OPCs are capable to functionally rescue nerve conduction after a demyelinating insult (Remaud et al., 2017), suggesting that exposure to any TH-disrupting chemicals could alter the regeneration of myelin, and may interfere with multiple sclerosis (MS) pathophysiology.

An unexplained increasing prevalence and incidence of MS occurred over the last 30 years (Magyari & Sorensen, 2019; Walton et al., 2020) with in addition a female/male sex ratio shifting from 2/1 to 3/1 for relapsing MS and even to 4/1 in some countries (Walton et al., 2020). In parallel, the quantity and diversity of chemicals used in our direct environment have considerably increased since 1970 (UNEP, 2012), leading to constant human exposure, from early development to aging. Among these industrial chemicals, PFAS are well established to interfere with TH signaling (Coperchini et al., 2020; Davidsen et al., 2022) and thus, constitute potential environmental cues that could disrupt TH-dependent processes as oligodendrogenesis and remyelination.

We investigated the adverse effects of these two perfluorooctanic acids on oligodendrogenesis and remyelination processes from cellular to behavioral levels. We demonstrated that in response to experimental demyelination PFOS, but not PFOA, impaired myelin regeneration by reducing the number of mature myelinating oligodendrocytes, together with functional consequences on sensory-motor behavior well-known to be linked to remyelination capacities. Our work combined *ex vivo* (organotypic cerebellar slice cultures) and *in vivo* approaches in two groups of vertebrates (mice and xenopus). This original approach allowed us to establish a comparative and evolutionary perspective of the action of PFAS on myelin physiology, opening to novel perspectives on the role of some environmental toxicants as a potential causative factor of some CNS demyelinating diseases such as MS.

## 2. Materials and methods

### 2.1. Animals

#### 2.1.1. **Mice**

C57/BL6 gestant females were purchased from Janvier (Le Genest-Saint-Isle, France) and kept in ventilated cages under a 12:12 h light-dark cycle in our animal facilities (agreement # A75-13-19. Experiments were conducted with respect to the European Union regulations and have been approved by the ethical committee of the French Ministry of Higher Education and Research (approval number Ce5/2010/025) (APAFIS #6269).

*In vivo* exposure to either PFOS or PFOA was induced by giving dams drinking water containing 20mg/L either PFOA or PFOS during a period corresponding to E15-p21 for the progeny. The bottle was renewed every three days. All experiments were approved by the ComEth ethical board (Project number APAFIS #21591-2019021815565069v7) and performed in strict accordance with European Directive 2010/63/EU.

#### 2.1.2. Xenopus

All experiments were performed on stage 48 to 50 Xenopus laevis tadpoles staged according to Nieuwkoop and Faber normal tables (Nieuwkoop PD, Faber J., s. d.)(. Tadpoles were obtained by natural mating of pairs of adults, selected from our colony of either transgenic Tg(*Mbp:GFP-ntr*) or wild type raised in our animal facility (agreement # A75-13-19). Handling of animals and functional tests have been previously detailed (Henriet et al., 2023). Animal care was in accordance with institutional and national guidelines. All animal procedures conformed to the European Community Council 1986 directive (86/609/EEC) as modified in 2010 (2010/603/UE) and have been approved by the ethical committee of the French Ministry of Higher Education and Research (APAFIS#5842-2016101312021965).

### 2.2. LC-MS/MS analysis

First, myelin of CTL, PFOS- and PFOA-exposed p21 mice were isolated following Percoll^®^ gradient cell separation from enzymatically dissociated brain tissue as previously described in (Moyon et al., 2021).

The internal standard M8PFOA was purchased from Wellington Laboratories (Canada). HPLC grade solvents were purchased from Merck and VWR.

PFOS and PFOA were extracted from myelin and serum using protein precipitation method. Briefly, 50µl of serum or 100µl of myelin in Percoll^®^ were supplemented with 7 volumes of acetonitrile and 10ng of M8PFOA internal standard. Samples were sonicated for 5min and proteins allowed to precipitate at 4°C for 1h. Samples were centrifuged at 20 000g for 20min at 4°C and the supernatant dried and resuspended in 50µl of methanol/water (1:1 v/v).

PFAS were quantified by LC-ESI/MS/MS using a Prominence UFLC (Shimadzu, Tokyo, Japan) and QTrap 4000 mass spectrometer (AB Sciex, Framingham, MA, USA) equipped with a turbo spray ion source (450°C) combined with an LC20AD HPLC system, a SIL-20AC autosampler (Shimadzu, Kyoto, Japan) and the Analyst 1.5 data acquisition system (AB Sciex, Framingham, MA, USA). PFAS were ionized in negative mode. Samples (4µl) were injected to a C18 Ascentis column (2.7µm, 2.1×150mm, Merck). Mobile phases consisted of (A) 2mM ammonium acetate and (B) acetonitrile. The gradient started with 30% of phase B and increased to 65% over 3min maintained for 1min and then increased to 100% over 3min. PFAS were detected using scheduled multiple reaction monitoring using sulfate fragmentation for PFOS 499<80 or neutral loss for PFOA 413<369 and M8PFOA 421<376. Myelin PFAS concentrations were estimated based on previously published rat brain composition (Norton & Poduslo, 1973): myelin weight was estimated at around 20-25% of total dry brain weight and brain water content at around 80% of total weight. Using these estimates and starting with mice brain total weight of 300mg, myelin weight was estimated at 13.5mg (300×0.2×0.225). Finally, PFAS weight was quantified in total Percoll^®^ fraction and divided by total myelin estimated weight.

### 2.3. Mouse cerebellar slice preparation

The procedure is described in the supporting information (see also (Thetiot et al., 2019)

### 2.4. Quantification of myelin on mouse cerebellar slice preparation

The procedure is described in the supporting information and (Ronzano et al., 2021)

### 2.5. Antibodies

List of antibodies is provided in the supporting information

### 2.6. Immunolabeling and quantification on brain section

The procedure is described in the supporting information

### 2.7. Quantification of GFP+ cells

GFP was detected directly by fluorescence in live Tg(*Mbp:GFP-ntr*) transgenic Xenopus embryos using an AZ100 Nikon Zoom Macroscope. The procedure is described in the supporting information (see also (Kaya et al., 2012) and (Henriet et al., 2023).

### 2.8. Behavioral testing

The tests have been previously described (Henriet et al., 2023). The procedure is described in the supporting information.

### 2.9. Statistical Analysis

We used Prism GraphPad software (GraphPad Prism version 8) for statistical analyses. Data presented are the mean ± SEM of number of GFP+ cells counted on at least 16 tadpoles per condition. For the analysis of two groups, an unpaired two-tailed Student t-test or a Mann-Whitney test were applied. For more than two group analyses, a one-way ANOVA with Tukey’s multiple comparison test or a Kruskal-Wallis with Dunn’s multiple comparisons test were applied. Statistical significance was defined as: *p < 0.05, **p < 0.01, and ***p < 0.001.

## 3. Results

### 3.1. In vivo accumulation into the myelin sheath of PFOS, and to a lesser extent of PFOA

PFAS are lipo-soluble compounds that accumulate into adipose tissue and other lipid rich-structures (Lee et al., 2017; Mamsen et al., 2019). Knowing the high lipid content of myelin sheath, oligodendroglial membrane wrapped around long projecting axons, we first interrogated whether a perinatal exposure to PFAS could accumulate into the pup’s myelin sheath. To test this possibility, either perfluorooctane sulfonate (PFOS) or perfluorooctanoic acid (PFOA) (20mg/L, each) was added to the drinking water of dams, during gestation from embryonic day 15 (E15) and during lactation till weaning, i.e., postnatal day 21 (p21) (**Fig. 1A**). At the end of PFAS exposure, similar level of either PFOS or PFOA were detected in the plasma of mothers (3.5±0.72 and 4.1±0.18 µg/ml, respectively; **Fig. 1B**) and of pups (1.63±0.19 and 0.86±0.13 µg/ml, respectively; **Fig. 1D**), indicating that both PFAS was similarly absorbed from the drinking water to the mother plasma and from the mother’s milk to the pup’s serum. On p21 myelin was bulk-purified from dams and pups’ brain by centrifugation on a Percoll^®^ gradient and collected myelin pellets were analyzed by LC-mass spectrometry. Both PFAS were recovered in the mother’s myelin fraction and 4.7 less for PFOS and 10 time less for PFOA in the pups’ myelin (**Fig. 1C, E**; and **supplementary Table 1**). Despite equivalent plasmatic levels, PFOS accumulated 133 time more than PFOA into the pups’ myelin (2.80±0.37 vs 0.021±0.003 ng/g tissue; Mann-Whitney test, p=0.0002), (**Fig. 1E**). Of note, low levels of both PFOS and PFOA were detected in the myelin of control pups, i.e., pups in which mother had been drinking normal tap water (**Fig. 1E**). To find the potential source of PFAS contamination in control animals we assayed the food pellets, litter and water. While hardly detectable in food pellets and litter, definite amount was present in the tab water (**supplementary Table 1**).

**Figure 1:**
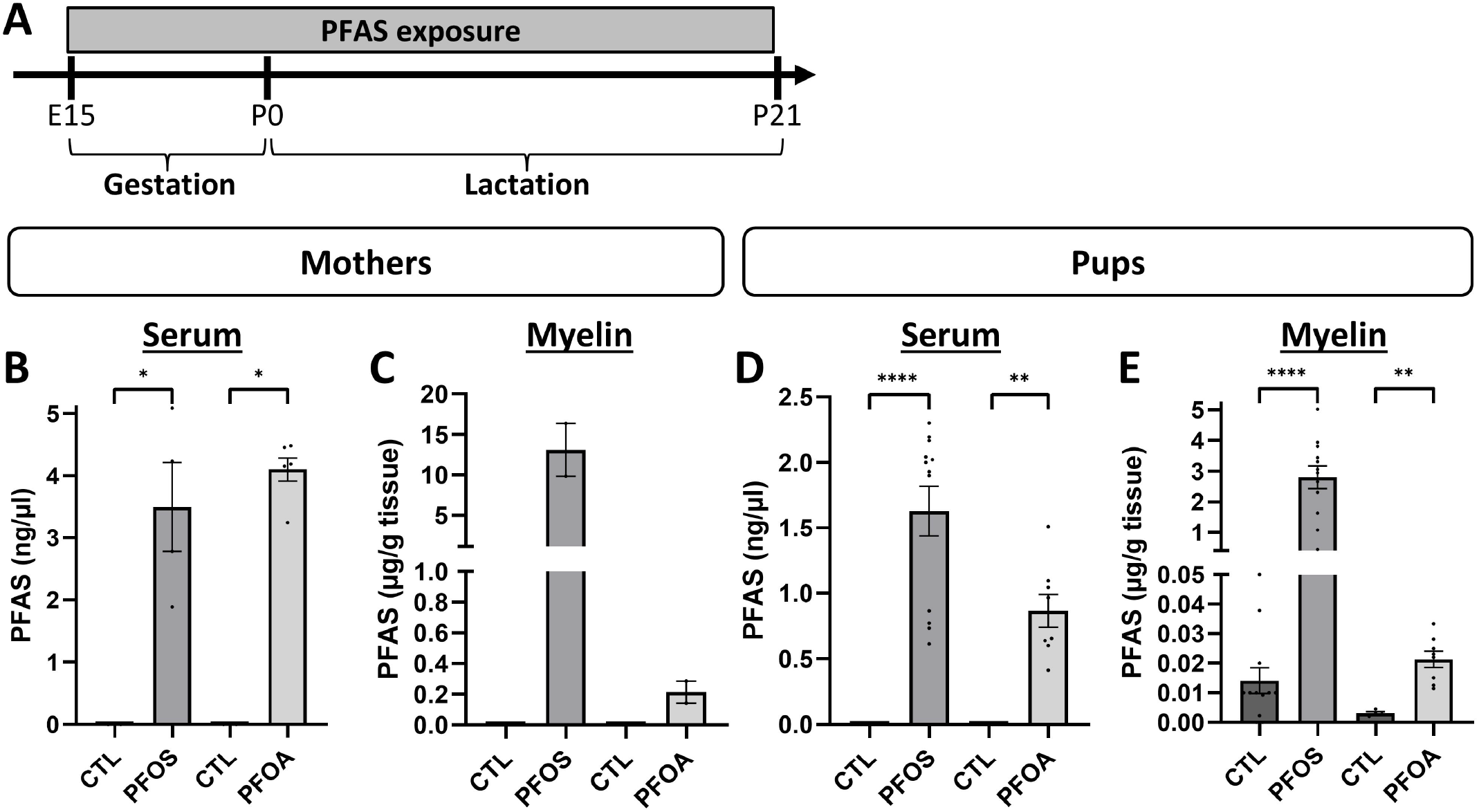
Accumulation of PFAS into pups’ myelin fraction. A) Flow chart of PFAS exposure (20mg/L) of adult female mice via the drinking water from E15 of gestation and during the lactation period. At 3 weeks (p21), dams and pups were euthanized, brains dissected out and myelin bulk fraction purified on a Percoll^®^ gradient. B-E) At p21, PFAS were assayed by LC-MS/MS in the serum of dams and pups (B, D) and corresponding myelin fractions (C, E). Although similar levels of either PFOS or PFOA passed from the drinking water into the mother serum and from the mother milk into the pups’ serum, 133 time more PFOS was detected into the pup’s myelin fraction compared to PFOA. Note that low level of both PFOS and PFOA were detected into the myelin of control animals (E), which is explained by the presence of these PFAS into the normal drinking water given to control dams (see supplemental table 1).

### 3.2. Exposure to PFOS, but not PFOA, altered oligodendrogenesis and myelination

We then examined whether the perinatal exposure to PFOS or PFOA affected the generation of new OPCs derived from the p21 dorsal SVZ niche by immunohistochemistry using an antibody directed against the oligodendroglial transcription factor OLIG2 (**Fig. 2A-D**). We observed that PFOS, but not PFOA significantly increased the density of OLIG2+ oligodendrocyte precursor cells (OPCs) (Dunn’s test, 0.027≤p≥0.04; **Fig. 2E**). In addition, the density of oligodendroglia lineage population at varying maturity was assessed in the corpus callosum above the dorsal SVZ by double-immunostaining with anti-OLIG2 to label all oligodendrocyte-lineage cells and a marker of mature oligodendrocytes (APC, recognized by CC-1 mAb) (**Fig. 2F-I**). We calculated ratios of OLIG2+/CC1+ co-expressing mature oligodendrocytes vs. immature OPCs expressing OLIG2 alone (**Fig. 2J**). About 70% of the OLIG2+ oligodendroglial cells also expressed CC1 in control mice. In PFOS-exposed animals, mature OLIG2+/CC1+ oligodendrocyte density decreased significantly (57%) in favor of an increase in the density of immature OPCs in the corpus callosum (Dunn’s test, p=0.006). The decrease in density of mature oligodendrocyte translated in a significant reduction of PLP myelin immuno-staining (Proteolipid Protein, PLP, the major myelin protein; **Fig. 2K-M**) (20% reduction, Dunn’s test, p=0.004) in the corpus callosum of PFOS-treated brains (**Fig. 2N**). Of note, on PFOA-exposed pups no significant effect was detected on oligodendrocyte differentiation (Dunns’ test, p>0.05; **Fig. 2E, J, N**).

**Figure 2:**
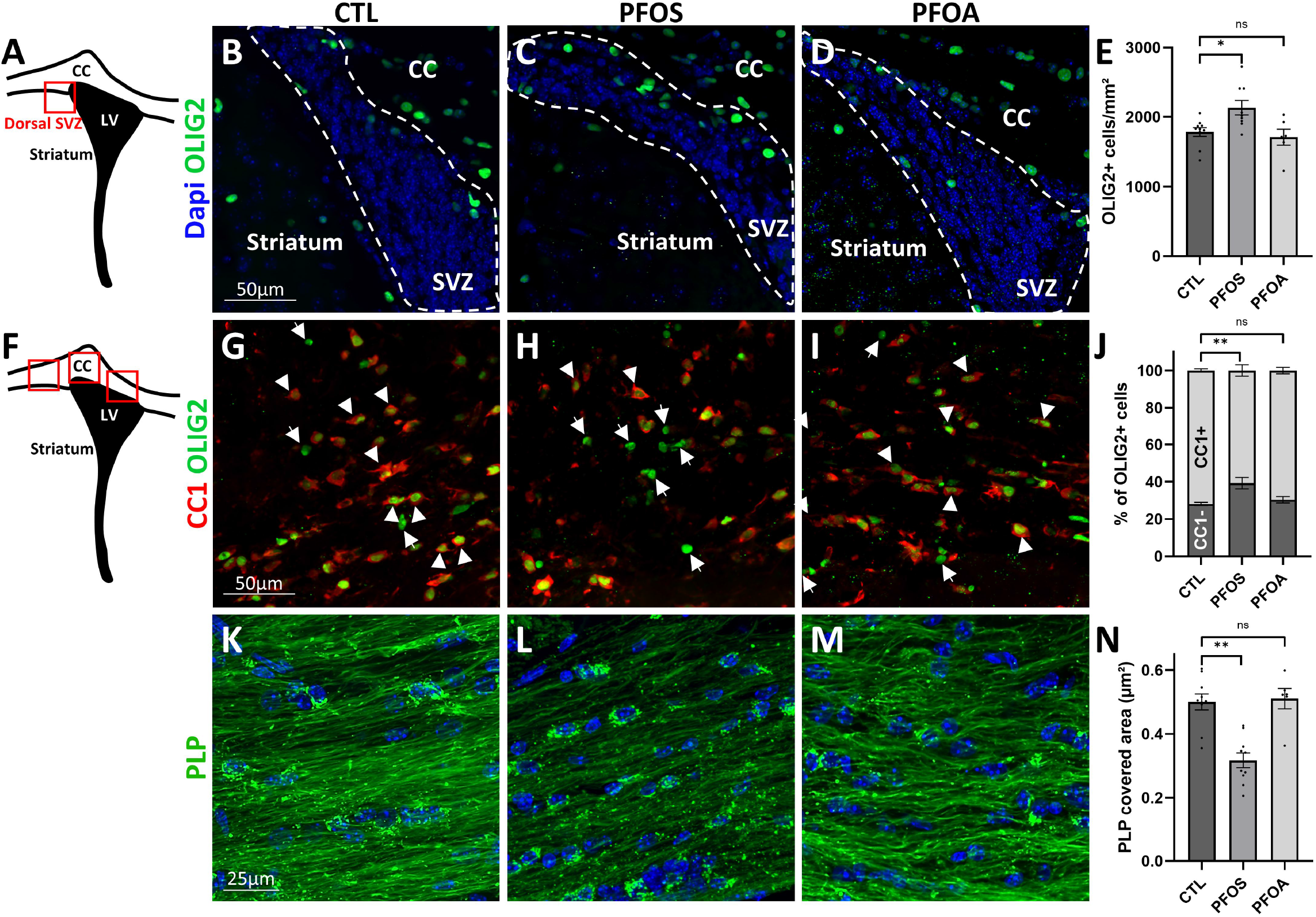
Perinatal exposure to PFOS, but not PFOA, increased the generation of SVZ-derived oligodendrocyte precursor cells (OPCs), which were not able to maturate into myelin-forming oligodendrocytes. A, F) Schematic drawing showing on coronal sections of p21 pups level illustrated in B-D for subventricular zone and in G-I and K-M for corpus callosum exposed to either PFOS, PFOA or control (CTL). B-E) section at the level of the SVZ immunostained for OLIG2 (green in B-D), at the level of the corpus callosum doubly stained for OLIG2 (green) and CC1 (red in G-I), and with anti-PLP to stain myelinated fibers (K-M) above the lateral ventricles. Note for PFOS exposed animals the increased density of OPC-OLIG2+ cells (C, E) contrasting with the lower ratio of mature oligodendrocytes OLIG2+/CC1+ (H, I) and the lower myelin staining (PLP+) in (L, N). In contrast, in brain sections of PFOA exposed animals, no significative differences compared to controls were observed. Nuclei are counterstained with DAPI; CC= corpus callosum, SVZ= subventricular zone, LV= lateral ventricles. Scale bar : B, C, D, F, G, H = 50µm; K, L, M = 25µm

### 3.3. PFOS altered remyelination in organotypic mouse cerebellar slices

Having shown the bioaccumulation of PFAS into the myelin sheath and that PFOS exposure impaired the generation of mature oligodendrocytes, we questioned whether PFAS affected remyelination. We used organotypic cerebellar slices from postnatal wild-type mice to better analyze the cellular effects of PFAS on remyelination *ex vivo*. Cerebellar slices from P9 wild type were cultured as described (Thetiot et al., 2019) (**Fig. 3A**). After 6 days in culture, cerebellar slices were transiently demyelinated for 15h using lysophosphatidylcholine (LPC). After LPC removal, endogenous remyelination of Purkinje cell axons occurred and slices were incubated for 4 days with decreasing concentrations of either PFOA (10^-7^ to 10^-9^M) or PFOS (10^-6^ to 10^-11^M). The extent of remyelination was measured by double -labeling of myelin and Purkinje axons using anti-PLP and anti-Calbindin antibodies, respectively (**Fig. 3B-G)**. The remyelination index was measured using an ImageJ macro language that allows a fast and an unbiased automated quantification of cerebellar myelinated axons (Ronzano et al., 2021). We observed that PFOS at concentration between 10^-8^ to 10^-10^ M, inhibited remyelination by 25-30% compared to control (Dunn’s test: 0.001≤p≥0.02), but not for the highest (10^-6^ or 10^-7^M) nor the lowest (10^-11^ M) concentration (**Fig. 3H).** This U shape dose-response curve is a characteristic of molecules acting on nuclear receptors (Vandenberg et al., 2012). Knowing the important role of thyroid hormone in oligodendrogenesis and myelin formation (Bernal, 2000; Remaud et al., 2017; Zorrilla Veloz et al., 2022), we first verified that in our *ex vivo* cerebellar culture explant demyelinated model, remyelination was strongly dependent on thyroid hormone signaling, increased by addition of T3 (10nM) in the culture medium and inhibited by addition of NH-3 a potent thyroid hormone receptor (THR) antagonist (**Supplementary Fig. 1**). (NH-3 is a derivative of the selective thyromimetic GC-1, which inhibits binding of thyroid hormones to their receptor) (Lim et al., 2002). We then interrogated whether the effect of PFOS could be reversed by T3 addition. Indeed, when slices were exposed to the most deleterious dose of PFOS (10^-8^M), addition of T3 (10nM) allowed to return the levels of remyelination close to the control (**Fig. 3H**), strongly suggesting that PFOS negative effect on remyelination could involve modulation of TH action. In contrast to data generated with PFOS, no significative effect of PFOA exposure was observed (**Fig. 3I;** Kruskal-Wallis test, p>0.05).

**Figure 3:**
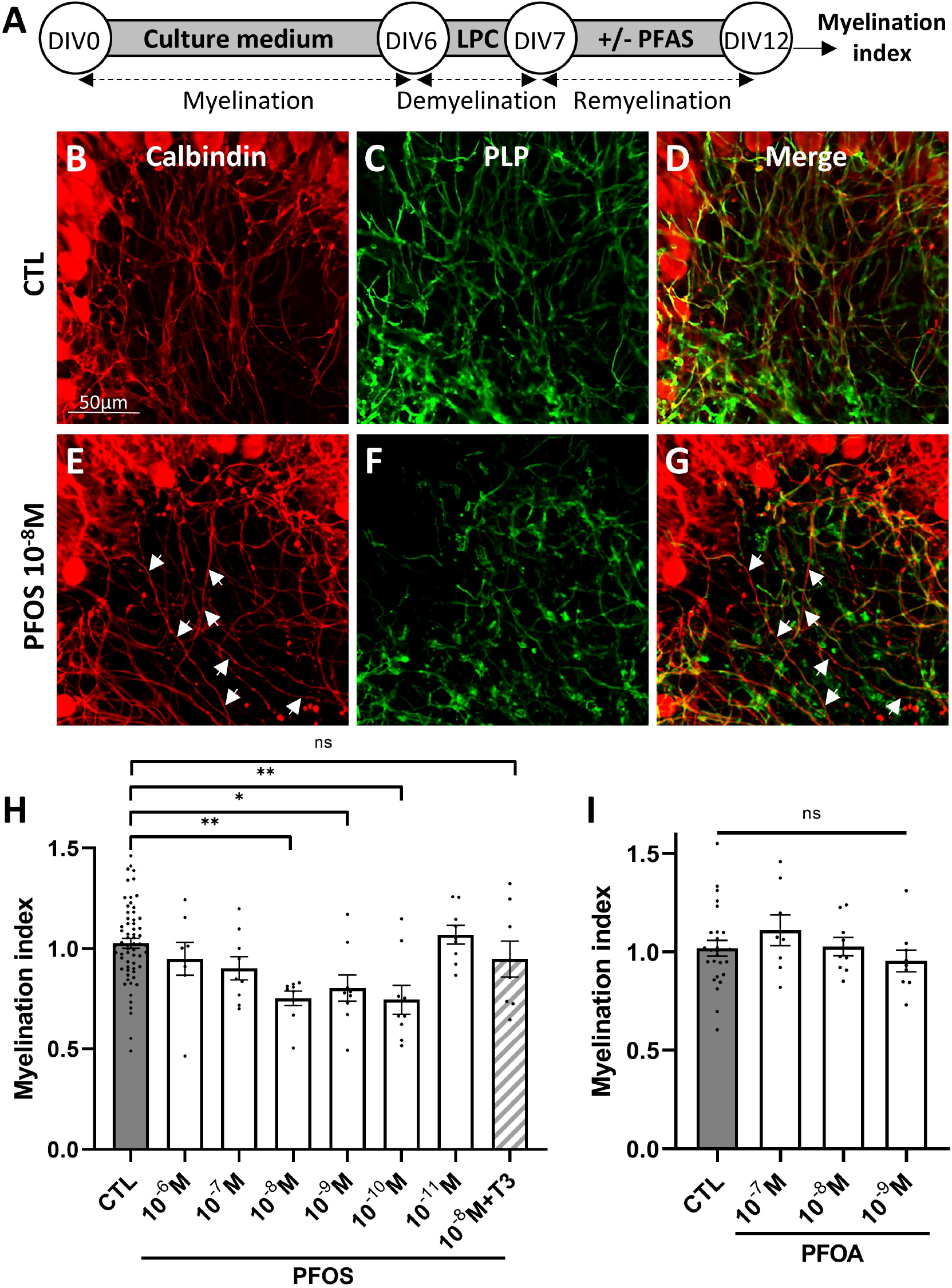
*Ex vivo* PFOS exposure inhibited remyelination. A) Flow chart showing the sequence of lysophosphatidylcholine-induced transient demyelination (from DIV6 to DIV7) of cerebellar slices followed by exposure to PFAS during the endogenous remyelination period, from DIV7 to DIV12. B-D) Cerebellar slices from P9 mice at 12 DIV doubly immunostained for calbindin (Purkinje cell bodies and axons in red in B, D, E, G) and PLP, (myelinated axons in green in C, D, F, G); note the decreased PLP+ myelin staining in slices exposed to PFOS, while Purkinje cells and axons appeared unaffected (E) compared to controls (B); arrow head in E and G point to axons not ensheathed by myelin. (H, I) Dose response curve of either PFOS (H) or PFOA (I) exposure on the myelin index measured as a ratio between the calbindin (axon)and PLP (myelin) stainings. Note the non-monotonic dose response of PFOS and the reversal of PFOS inhibitory effect on remyelination by addition of T3 into the culture medium (hachured column in H). In contrast PFOA (I) did not affect remyelination. Scale bar: B-G= 50µm.

### 3.4. Functional consequences of PFAS-induced impaired remyelination in Xenopus

We then interrogated the functional consequence of PFOS bioaccumulation. In a first set of experiments we took advantage of our transgenic Tg(*Mbp:GFP-ntr*) Xenopus line to test the possible deleterious effect of PFAS on myelin biology and more specifically the endogenous repair potential. In this transgenic line, the GFP reporter is expressed specifically and selectively in myelin forming oligodendrocytes. In addition, due to the nitroreductase (NTR) transgene -fused to the GFP reporter-myelinating oligodendrocytes were ablated following addition of metronidazole (MTZ) into the swimming water (**Fig. 4A-F**). The hydroxylamine product of the NTR-induced reduction of the NO2 moiety of MTZ is highly toxic, leading to an oligodendrocyte cell death. Depending on the concentration and duration of MTZ exposure the extent of oligodendrocytes ablation is more or less severe and was best quantified in the optic nerve. In stage 48-50 the number of GFP+ oligodendrocyte was very stable and reproducible (16.54 ± 0.64; n=237) and was severely decreased at the end of the demyelination treatment −10 days exposure to MTZ (10mM)-(4.25 ± 0.92; n=176)(**Fig. 4C-D**). Following this demyelination period when tadpoles were returned to normal water spontaneous recovery occurred. After 3 days the number of GFP+ oligodendrocytes was 13.98 ± 0.57 (n=44) (**Fig. 4E, G**) and after 8 days recovery was complete or almost complete. We have previously shown that the number of GFP+ oligodendrocytes per optic nerve was a faithful and reliable index of myelin content, whether at the end of the demyelination period as well as during remyelination and that this model was sensitive enough to identify among a panel of compounds added into the swimming water the ones that had the potential to promote remyelination(Kaya et al., 2012; Mannioui et al., 2018). To examine the effect of PFAS on remyelination, at the end of the MTZ-induced demyelination tadpoles were exposed during the 3 days period of recovery to decreasing doses ranging between 10^-5^ M and 10^-11^ M of either PFOA or PFOS (**Fig. 4A, F, G, H**). Compared to control levels, i.e., level of spontaneous recovery, no significative effect on the number of oligodendrocytes was observed at any of PFOA concentrations tested (Kruskal-Wallis test, p>0.05; **Fig. 4H)**. In contrast, PFOS induced a maximal 37.2% inhibition of recovery for a concentration of 10nM (Dunn’s test: p<0.0001; **Fig. 4G)**. The inhibitory effect was dose dependent with a U-shape curve, with no effect for the highest (10^-5^ M) or lowest concentration (10^-11^ M) and a maximum effect for 10^-8^ M.

**Figure 4:**
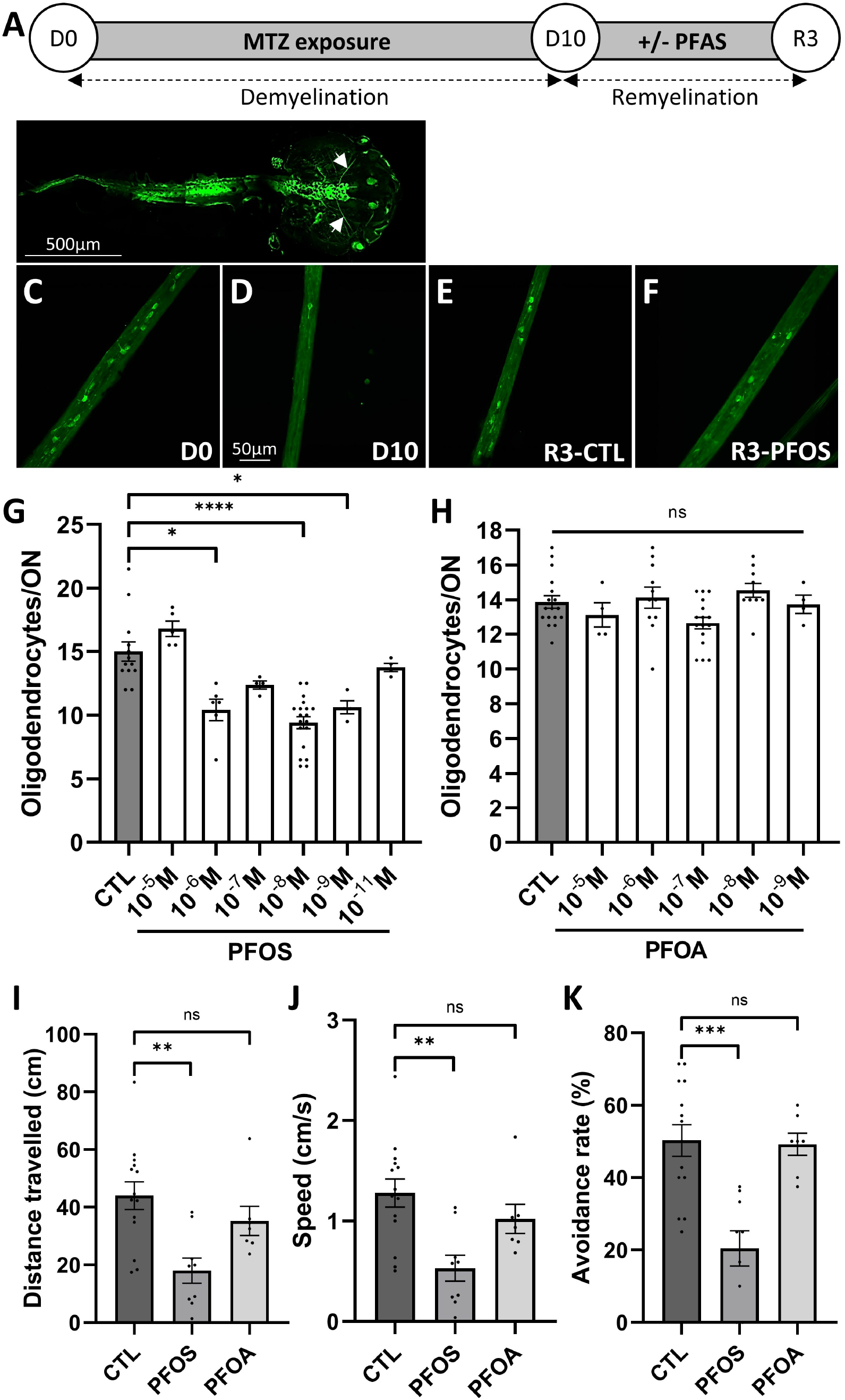
*In vivo* PFOS inhibited remyelination. (A) Flow chart showing the sequence of events tested. (B) Whole mount of Tg(*Mbp:GFP-ntr*) Xenopus laevis (stage 50 tadpoles) showing GFP expression in the brain, optic nerve (white arrows) and spinal cord. At this magnification GFP+ oligodendrocyte cell bodies are not visible. (C-F) Higher magnification of Tg(*Mbp:GFP-ntr*) Xenopus laevis optic nerve before (D0)(C), at the end of metronidazole-induced demyelination (D10)(D) and after 3 days (R3) of spontaneous recovery in control (E) or PFOS exposed tadpoles (F); scale bar in B= 500µm in C-F = 50 µm. (G, H) Dose response of remyelination after addition for 3 days of either PFOS (G) or PFOA (H). Remyelination was evaluated by counting the number of GFP+ oligodendrocytes per optic nerve. Note the non-monotonic inhibition of remyelination following PFOS exposure, while PFOA had no effect. (I-K) PFOS inhibition of remyelination affected tadpoles’ behavior as assayed by distance traveled (I), speed of swimming (J) and visual avoidance of a virtual collision (K) contrasting with the absence of effect of PFOA.

We questioned the functional consequences of the decrease in the number of mature myelin forming oligodendrocytes. To evaluate the motor functions of the tadpole we measured the distance traveled for a given period of time and the speed of swimming. After 10 days in MTZ (10mM) demyelinated animals swam a shorter distance than before demyelination: 57.3 ± 3.12 cm vs 10.13 ± 1.39 cm at D0 and D10, respectively (m ± SEM, n=63 (D0), n=36 (D10); Dunn’s test: p= <0.0001). Similarly, the average speed of swimming of Tg(*Mbp:GFP-ntr*) tadpoles (1.67 ± 0.09 cm.s^-1^) was significantly decreased at the end of the demyelination treatment (0.30 ± 0.04 cm.s^-1^) (mean ± SEM, n=63 (D0), n=36 (D10); Dunn’s test: p= <0.0001).

At R3, the nearly complete recovery in the number of GFP+ cells measured in the optic nerve paralleled an improvement in the average speed and the distance traveled over a period of 30s (distance: 44 ± 4.8 cm; speed: 1.28 ± 0.14 cm.s^-1^; n=14). In contrast, tadpoles exposed to PFOS (10nM) did not recover (distance: 18.02 ± 4.36 cm; speed: 0.53 ± 0.13 cm.s^-1^; n=9; Dunn’s test: p=0.004) (**Fig. 4 I, J)**.

Since the level of demyelination was evaluated by the number of mature GFP+ oligodendrocytes per optic nerve, we reasoned that a demyelination, or failure of remyelination of optic nerve axons must translate by a loss of vision, like optic neuritis in human. To test whether a demyelination of optic nerve axons translated in tadpoles into a vision loss we developed an index of visual avoidance of a virtual collision (Henriet et al., 2023). At the end of the demyelination period, tadpoles had lost the capability to avoid the threatening stimulus represented by the virtual collision with the black dot. The avoidance index measured at D0 significantly decreased after 10 days of demyelination from 69.76 ± 1.4% to 22.27 ± 2.89%; n=63 (D0), n=36 (D10); (mean ± SEM; Dunn’s test: p=<0.0001) (**Fig. 4K**). Three days after MTZ exposure was stopped, control animals recovered rapidly with avoidance index of 50.28 ± 4.35% (n= 14) while PFOS treated tadpoles did not recover (avoidance: 20.43 + 4.88%; n=9; Dunn’s test: p=0.0007) (**Fig. 4K**). In contrast to PFOS, and similar to our observations in mouse cerebellar slices, in xenopus PFOA had no noticeable effect on cellular remyelination (**Fig. 4H)** and did not alter spontaneous functional repair (**Fig. 4I-K**).

## 4. Discussion

### 4.1. Comparative analysis of PFOA and PFOS effects

PFOA and PFOS were given at 20mg/L via drinking water to gestant and lactating mothers and both compounds were detected at similar levels in the serum of p21 pups by LC-MS/MS, showing placental transfer of PFAS as previously reported (Chen et al., 2017). The high concentration of PFAS in blood could be also explained by the high affinity of PFAS for plasma albumin (Guy WS, s. d.). Furthermore, although prenatal exposure to PFOS and PFOA has been usually associated with lower birth weight in humans (Fei et al., 2007) and rodents (Lau et al., 2004, 2006)(for PFOS (2–20 mg/kg) and PFOA (1–40 mg/kg), we did not observe any significant modifications of birth weight and postnatal survival in both PFAS-treated groups (11.21 ± 0.32g and 10.98 ± 0.41g for PFOS and PFOA, respectively) compared to controls pups (11.72 ± 0.47g), suggesting that *in vivo* PFAS exposure at 20mg/L has limited adverse health effects on mouse perinatal development. A large part of the literature is focused on the association of PFAS with protein-rich tissues (liver and blood) rather than lipids (Jones et al., 2003). In human, PFAS are mostly detected in lung and liver and to a lesser degree in the central nervous system (Mamsen et al., 2019). Cerebral barriers may limit PFAS entry into the brain although blood-brain-barrier disruption could be a mechanism facilitating PFOS entry in brain (35), as occurring in inflammatory CNS diseases. Interestingly, the levels and patterns of PFAS have been reported in several brain regions (i.e., notably hypothalamus, striatum, cerebellum, cortex of polar bears (Greaves et al., 2013) but, to our knowledge, accumulation of PFAS in the white matter has never been clearly identified. Our study emphasizes not only a preferential PFAS accumulation into the myelin, but also a higher occurrence of PFOS than PFOA.

The differential accumulation into the myelin sheath of PFOS compared to PFOA may result from different, not exclusive factors: i) PFOS is a higher hydrophobic compound with its SO3H function that may facilitate its incorporation into the lipid-rich sheaths; ii) although the existing literature shows that PFAS biotransformation is minimal or absent (Vanden Heuvel et al., 1991), PFOA exhibits much faster depuration than PFOS (Benskin et al., 2009; Hassell et al., 2020) and PFOS has a greater half-life in tissues than PFOA (Li et al., 2018). In addition, competition for TH binding sites on serum transport proteins, such as transthyretin (TTR), which binds and distributes TH to brain cells *via* the cerebrospinal fluid may facilitate PFAS brain entry, notably in brain cells closed to the ventricles (Vancamp et al., 2019). In this context, it is noteworthy that PFOS has a higher TTR-binding potency compared to PFOA (Weiss et al., 2009), which may favor PFOS delivery to neural cells.

### 4.2. The myelinotoxic and gliotoxic mechanisms of PFOS

We demonstrated that PFOS accumulated into the myelin fraction of p21 pups. Further work is needed to determine whether accumulation of PFAS into the myelin sheath could alter myelin functions. In particular, a lipidome and proteome map of myelin membranes combined to ultrastructure analysis would allow to better understand the molecular anatomy of the potential PFOS-induced myelin deterioration.

We also showed *in vivo* that PFOS impaired oligodendrocyte lineage formation, notably the differentiation of mature myelinating oligodendrocytes from neural stem cells localized within the SVZ. The pool of immature precursors is increased at the expense of differentiated CC1+ and PLP+ oligodendrocytes within the corpus callosum, the white matter just above the SVZ where newly generated SVZ-derived OPCs migrate (Remaud et al., 2017). A similar phenotype has been described in several genetic and pharmacological models deficient for the TH signaling (Gothié et al., 2017; Luongo et al., 2021; Remaud et al., 2017; Vancamp et al., 2019), strongly suggesting that these PFOS-related effects involved modulation of TH action. Furthermore, myelin deposition is a well-established T3-dependent process in all vertebrates (Barres et al., 1994; Billon et al., 2001). Accordingly, addition of T3 rescued the negative effect of PFOS (10^-8^ M) on endogenous myelin repair *ex vivo*. The ability of PFAS, notably PFOS, to disturb TH biosynthesis and metabolism is well documented in animal models (Yu et al., 2009) and humans (Boas et al., 2012; Melzer et al., 2010), making PFOS a TH-disrupting chemical.

The activity of PFOS on oligodendrogenesis and myelination in the CNS could also be linked to its interaction with many others nuclear hormone receptors (i.e., PPAR, estrogen or androgen pathways (Villeneuve et al., 2023) well-known to be modulated by PFOS in peripheral tissues (Du et al., 2013; Wan et al., 2012) and to regulate myelin homeostasis (Hussain et al., 2013; Montani et al., 2018; Taylor et al., 2010).

### 4.3. Is there a link between PFOS exposure during the perinatal period and the prevalence of multiple sclerosis?

We have previously demonstrated that the perinatal period is a critical developmental window sensitive to endocrine disruption that could lead to neurogliogenic and behavioral permanent alterations in the adult (Vancamp et al., 2022, 2023). This work is in line with “the Developmental Origins of Health and Disease” (DOHaD) theory involving that early exposure to endocrine disrupting chemicals could induce irreversible damage later in life. It has been suggested that Persistent Organic Pollutants (POP) exposure, which includes PFAS, during the perinatal period may play a role in several neurological disorders such as ADHD, Autism Spectrum Disorder, Alzheimer’s Disease, Parkinson’s Disease. A link with myelin disorders has not been hypothesized (for review see (Grova et al., 2019)and to our knowledge, PFAS early exposure has not been established as a risk factor for MS, yet. A recent study, however, pointed out that POP exposure, including PFAS, decreased some myelination related genes (Yadav et al., 2022).

Aside from genetic factors, which only explain a fraction of the disease risk, and Epstein-Barr virus (EBV) infection, which increases by approximately 30 fold the risk of developing MS (Bjornevik et al., 2022), some environmental MS risk factors, - such as smoking, low vitamin D levels caused by insufficient sun exposure and/or dietary intake, obesity during adolescence-, have been identified and might participate to the increased incidence of the disease detected during the last decades, with odds ratio not exceeding 2. Increased exposure to EDC, notably PFOS, during the last decades is striking, and our work highlights their accumulation in CNS myelin on the one hand, their impact on oligodendroglial fate during development and on myelin integrity in the adult CNS, on the other hand. Therefore, although caution is needed because our data on PFAS accumulation are derived from mouse brains, PFOS exposure – and more largely EDC - might represent a novel and major risk factor for MS development, a hypothesis in line with inside-out pathogenic concept for MS, where initial myelin alterations can trigger an autoimmune CNS pathology (Luchicchi et al., 2021). Along this line, it is conceivable that PFOS accumulation into the myelin sheath, by modifying the lipid composition or by intercalating within the lipid bilayer, might alter the myelin biophysical properties, fragilize the membrane and its stability. EDCs are chemicals that either mimic or block endogenous hormones and thus disrupt the normal hormone homeostasis. So, there may be a possibility to counteract their deleterious effects by using agonists of the pathway they are perturbating. But in our opinion, the future way to go is prevention. Our hope is that our pioneer work will lead many other colleagues towards this avenue of research and therefore create a pressure on environmental protection authorities, not only to prohibit the manufacturing of these molecules, but also to develop systems to protect the population from the existing persistent pollutants and promote the depollution of contaminated soils and waters.

## Acknowledgements

We thank Drs Francine Acher, Dominique Padovani and Erwan Galardon for precious advises on different chemical properties between PFOS and PFOA, Drs Allan Tobin and Vincent Laudet for careful reading and helpful advices on the manuscript and David Akbar and ICM-QUANT imaging facility for help in generating micrograph illustrations. We also thank Fabien Uridat and Stéphane Sosinsky (UMR 7221) for excellent animal care. We thank Pamela Lein for supplying the NH3 (Walter et al., 2021).

## Authors’ contributions

PJ, LB, EM, MSA, ML, BZ and SR performed the experiments and analyzed the data; SR and BZ wrote the manuscript; JBF, BD, BS and CL were involved in revising the manuscript critically for important intellectual content and made substantial contributions to interpretation of data. BZ and SR conceived the study and supervised experiments. All authors read and approved the final manuscript.

## Classification

Biological Sciences: Systems Neuroscience; Cellular and Molecular Neuroscience;

## 8. Supporting information

### 8.1. Antibodies

Mouse mAb anti-Calbindin (IgG1, Sigma C9848, 1:800), rat mAb anti-PLP (culture supernatant 1:20; kindly provided by Dr. K. Ikenaka, Okasaki, Japan), chicken anti-GFP (1:500; Millipore); rabbit anti-Olig2 Ab (diluted 1:300, Millipore, AB15328), and a mouse IgG2b anti– adenomatous polyposis coli (APC, clone CC1; diluted 1:300, Calbiochem). Corresponding Alexa Fluor secondary antibodies were from Invitrogen (Thermo-Fisher) and all were used at a dilution of 1:600).

### 8.2. Mouse cerebellar slice preparation

Mouse cerebella from p9 animals were dissected in ice cold Gey’s balanced salt solution complemented with 4.5 mg/ml d-Glucose and penicillin-streptomycin (100 IU/mL, Thermo Fisher Scientific). They were cut into 300 μm parasagittal slices using a McIlwain tissue chopper and the slices placed on Millicell membrane (3–4 slices per membrane, 2 membranes per animal, 0.4 μm Millicell, Merck Millipore) in 50% BME (Thermo Fisher Scientific), 25% Earle’s Balanced Salt Solution (Sigma), 25% heat-inactivated horse serum (Thermo Fisher Scientific), supplemented with GlutaMax (2 mM, Thermo Fisher Scientific), penicillin–streptomycin (100 IU/mL, Thermo Fisher Scientific), and d-Glucose (4.5 mg/ml; Sigma). Cultures were maintained at 37 °C under 5% CO_2_ and medium changed every two to three days. At DIV6, demyelination was induced in cerebellar slices by treatment with lysophosphatidylcholine (LPC 50 mg/ml) for 15h. After 3 washes in 1X PBS, slices were incubated in a fresh culture medium for 4h then in medium supplemented with EDCs. At 12 DIV, cerebellar slices were fixed with 4% paraformaldehyde for 10 min and washed 3 times in 1X PBS. Slices were then blocked with 10% normal goat serum and doubly immunostained with a combination of anti-PLP and anti-Calbindin After 3 washes, cerebellar slices were incubated with corresponding Alexa-conjugated fluorescent secondary antibodies and mounted on glass slides with DAPI Fluoromount-G (InVitrogen).

### 8.3. Quantification of myelin on mouse cerebellar slice preparation

To analyze the effect of PFAS treatments on remyelination *ex vivo,* three folia per cerebellar slices were acquired per condition for each animal (2400 × 2400 pixels11 z-series with a z-step of 0.375 µm). The myelination index was calculated semi-automatically using a custom written script on ImajeJ. Briefly, a region of interest including Purkinje cells axon (excluding soma and white matter tracks) was first selected. A mask for axonal area (Calbindin signal) and a mask for myelinated axonal area (PLP signal overlapping with Calbindin signal) were then generated, and the myelination index was calculated from the quotient of the area of the two respective masks (myelin/axon). Myelination indexes of the five images were averaged to give the mean myelination index per animal for each condition.

### 8.4. Immunohistochemistry and quantification on brain section

Brains were fixed in 4% paraformaldehyde in PBS overnight at 4°C, cryoprotected in 30% sucrose and embedded in OCT (Sakura Finetek, the Netherlands) before being frozen and stored at −80°C. Coronal floating sections (20 µm) were made on a cryostat and stored at −20°C in cryoprotectant solution. P21 mice (n=10 CTL, n=10 PFOS and n=6 PFOA) were used for immunohistochemistry. Sections (n=3-4 per mouse) were incubated for 30 min in blocking solution 1% BSA (Sigma), 0.3% Triton X-100 and 10% donkey serum (Sigma) in 1X PBS at room temperature (RT), and then incubated with primary antibodies (anti-OLIG2, CC1 and anti-PLP) diluted in the same solution overnight at 4°C. After 3 washes (for 5 min) in 1X PBS at RT, sections were incubated with Alexa-conjugated fluorescent secondary antibodies (1% BSA, 0.3% Triton X-100 and 1% donkey serum in 1X PBS) for 2h at RT. Following incubation with DAPI for 5 min at RT, sections were covered with Prolong Gold antifade reagent (Invitrogen) and sealed with coverslips. Images were acquired using a Nikon confocal microscope under 400x magnification. The dorsal SVZ was imaged to analyze the density of OLIG2+ OPCs. To quantify OPCs vs mature oligodendrocytes and myelin in the corpus callosum, three images were acquired per section. Cell quantification was performed with FIJI software, using the cell counter plugin. The cell density (number/mm2) was calculated by counting the number of immuno-positive cells per area. For PLP, the labeling surface was automatically quantified and related to the area surface.

### 8.5. Quantification of GFP^+^ cells

GFP was detected directly by fluorescence in live Tg(*Mbp:GFP-ntr*) transgenic *Xenopus* embryos using an AZ100 Nikon Zoom Macroscope. The total number of GFP^+^ cells was counted in the optic nerve, from the emergence of the nerve (i.e., after the chiasm) to the retinal end. For stage 50 tadpoles the length of the optic nerve is on average 1700 µm ± 100 µm for a diameter of 50 µm. GFP^+^ cells were counted before (D0) and at the end of MTZ exposure (D10) and after being returned to normal water for 3 days (R3) or water containing the PFAS to be tested on the same embryos. Counts were performed independently by two researchers. Difference in numbers obtained by each researcher was below 10%. Data were compared to control untreated animals of the same developmental stage.

### 8.6. Behavioral testing

Tadpoles were tested in the morning before being fed. The setup consists of a CRT monitor (Dell Model #M570, 100-240 V, 60/50 Hz, 1.4 A, refresh rate used 60 Hz). The screen was covered with a 10mm diameter mask, adapted to a petri dish. Movement of tadpoles were recorded with a Dragonfly2 DR2-HIBW camera at 30 pfs and the Computar M3Z1228C-MP2/3” 12-36 mm Varifocal, Manual Iris Megapixel (C mount) lens. The video recording system used was FlyCapture2.

The setup was localized in a darkroom, light was turned off so that the only light perceived by tadpoles came from the screen. Each animal was tested separately in the Petri dish filled up to 1cm with MMR 0.1 X medium. Tadpoles were placed in the Petri dish and left to adapt to the screen-light for 5-10 s. Spontaneous swimming was recorded for 30 s and average speed for this period analyzed. If the animal was immobile at first it was touched with a plastic pipette to initiate movement, this first acceleration being excluded from analysis.

The virtual avoidance collision test was performed after all animals had been tested for spontaneous swimming behavior. A black dot (18 pixel = 8 mm on the screen) was presented on the screen, the experimenter targeted the eye of the tadpole by changing the direction and speed of the dot. On average, 5-6 tryouts were performed to assess visual avoidance. The virtual collision setup can be found on AK web site https://github.com/khakhalin/Xenopus-Behavior

Analysis of videos recordings was with Noldus Ethovision XT 11.5 software. For each experiment detection settings were calibrated. After tracking of the tadpole and the moving black dot the trajectories were individually verified and modified in case of swapping identity between tadpole and dot or in case of failure of automated detection. To determine visual avoidance several escape responses were analyzed and it was determined that a successful avoidance response corresponded to an acceleration swim of the tadpole >50 cm.s^-2^ and a change in direction (C-start) verified by the experimenter, initiated for a distance between the tadpole and the dot of 1-1.3 cm. Data are presented as an avoidance rate, i.e., the ratio of the number of encounters that resulted in a successful avoidance.

## 9. Legend to Supplementary Figure 1

### Remyelination is dependent on thyroid hormone signaling

Spontaneous remyelination of mouse cerebellar slices demyelinated by LPC is increased by addition of T3 (10nM) into the culture medium and inhibited by NH3 (5µM) a thyroid hormone receptor antagonist. Purkinje cells and axons were immunostained with anti-calbindin (A, C, D, F) and myelin sheath with anti-PLP (B, E). G) Myelination index was evaluated in all 4 conditions.

## 10. Supplementary Table 1

Level of PFOS and PFOA (assayed by LC-MS/MS) in serum, myelin and in the milieu (water, food pellet and litter).

## 11. Contact and competing interest information for all authors

The authors have no conflict of interest to declare relevant to this manuscript.

