## Supplemental Figure 1 and Table for "Deleterious functional consequences of perfluoroalkyl substances accumulation into the myelin sheath"

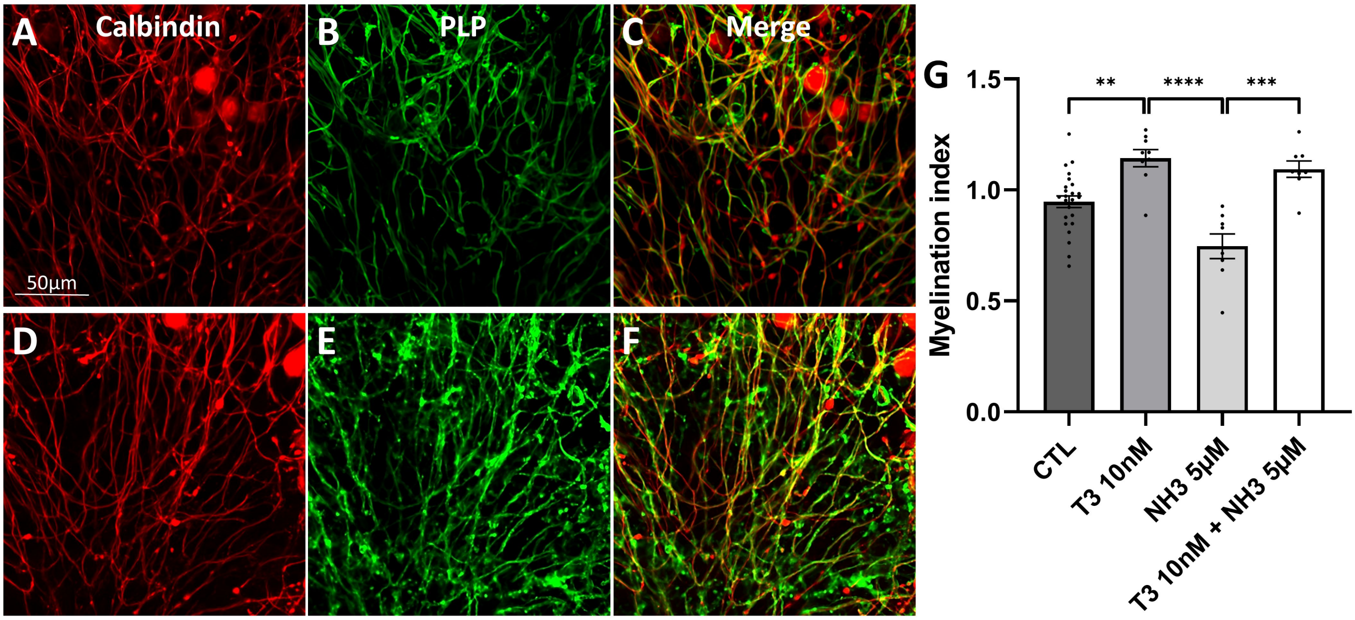

### Table 1

| | | P21 | | | Mothers | | |
|---|---|---|---|---|---|---|---|
| | | CTL | PFOS | PFOA | CTL | PFOS | PFOA |
| Water (ng/ul) | PFOS | | | | 0.0086 | 120.06 | 0.0832 |
| | PFOA | | | | 0.1 | 0.4653 | 137.4343 |
| Food (ng/g) | PFOS | | | | ND | | |
| | PFOA | | | | 0.0057 | | |
| Litter (ng/g) | PFOS | | | | 0.0146 | 12.3890 | 0.0227 |
| | PFOA | | | | 0.0083 | 1.3480 | 57.2746 |
| Serum (ng/µl) | PFOS | 0.0009 ± 0.0003 | 1.629 ± 0.191 | 0.0036 ± 0.0008 | 0.0004 ± 0.0001 | 3.496 ± 0.718 | 0.0122 ± 0.0032 |
| | PFOA | 0.0005 ± 0.0003 | 0.0036 ± 0.0008 | 0.8654 ± 0.0008 | 0.0004 ± 0.0001 | 0.0189 ± 0.0079 | 4.102 ± 0.184 |
| Myelin (ng/g tissue) | PFOS | 0.014 ± 0.004 | 2.81 ± 0.37 | 0.0044 ± 0.0016 | 0,0025 ± 0,0005 | 13,07 ± 3,27 | 0,04 ± 0,009 |
| | PFOA | 0.0031 ± 0.0005 | 0.0021 ± 0.0005 | 0.0213 ± 0.0027 | ND | 0,0017 ± 0,0001 | 0,21 ± 0,07 |
